# A genome-wide association study of mitochondrial DNA copy number in two population-based cohorts

**DOI:** 10.1101/372177

**Authors:** Anna L. Guyatt, Rebecca R. Brennan, Kimberley Burrows, Philip A. I. Guthrie, Raimondo Ascione, Susan Ring, Tom R. Gaunt, Angela Pyle, Heather J. Cordell, Debbie A. Lawlor, Patrick Chinnery, Gavin Hudson, Santiago Rodriguez

## Abstract

Mitochondrial DNA copy number (mtDNA CN) exhibits interindividual and intercellular variation, but few genome-wide association studies (GWAS) of directly assayed mtDNA CN exist.

We undertook a GWAS of qPCR-assayed mtDNA CN in the Avon Longitudinal Study of Parents and Children (ALSPAC), and the UK Blood Service (UKBS) cohort. After validating and harmonising data, 5461 ALSPAC mothers (16-43 years at mtDNA CN assay), and 1338 UKBS females (17-69 years) were included in a meta-analysis. Sensitivity analyses restricted to females with white cell-extracted DNA, and adjusted for estimated or assayed cell proportions. Associations were also explored in ALSPAC children, and UKBS males.

A neutrophil-associated locus approached genome-wide significance (rs709591 [*MED24*], β[SE] −0.084 [0.016], *p*=1.54e-07) in the main meta-analysis of adult females. This association was concordant in magnitude and direction in UKBS males and ALSPAC neonates. SNPs in and around *ABHD8* were associated with mtDNA CN in ALSPAC neonates (rs10424198, β[SE] 0.262 [0.034], *p*=1.40e-14), but not other study groups. In a meta-analysis of unrelated individuals (N=11253), we replicated a published association in *TFAM* β[SE] 0.046 [0.017], *p*=0.006), with an effect size much smaller than that observed in the replication analysis of a previous *in silico* GWAS.

In a hypothesis-generating GWAS, we confirm an association between *TFAM* and mtDNA CN, and present putative loci requiring replication in much larger samples. We discuss the limitations of our work, in terms of measurement error and cellular heterogeneity, and highlight the need for larger studies to better understand nuclear genomic control of mtDNA copy number.

## Introduction

Mitochondria are the cellular organelles responsible for producing adenosine triphosphate (ATP), a ubiquitous substrate required for metabolism. ATP is the final product of the series of redox reactions that are facilitated by the complexes of the respiratory chain (RC), located on the cristae, the folded inner membrane of mitochondria.

Mitochondria possess their own genome (mtDNA), an extra-nuclear, double-stranded, circular DNA molecule of ~16.6kb that is inherited maternally. Thirteen subunits contributing to complexes of the RC are encoded by mtDNA, and the entire mitochondrial genome is present at variable copy number in the cell. The relative copy number of mtDNA (mtDNA CN) may reflect differing energy requirements between cells: those from active tissues (e.g. liver, muscle, neuron) are observed to have higher mtDNA CNs compared to endothelial cells, which are comparatively quiescent.(Moyes et al. 1998; Xing et al. 2008; Dickinson et al. 2013)

Several nuclear genes are known to influence the regulation of mtDNA CN, and these are reviewed in detail elsewhere.(Rotig and Poulton 2009; Carling et al. 2011; Harvey et al. 2011; Dickinson et al. 2013) These include *POLG*(Oskoui et al. 2006; Bornstein et al. 2008; Rotig and Poulton 2009; Spinazzola et al. 2009; Tyynismaa et al. 2009; Carling et al. 2011; Chiaratti et al. 2011; Harvey et al. 2011; Venegas et al. 2011) and *POLG2,*(Tyynismaa et al. 2009; Carling et al. 2011; Harvey et al. 2011) the catalytic and accessory subunits of DNA polymerase-gamma, the principal enzyme implicated in mtDNA replication. Other regulators include *TFAM* (mitochondrial transcription factor A),(Choi et al. 2006; Belin et al. 2007; Curran et al. 2007; Alvarez et al. 2008; Carling et al. 2011; Cai et al. 2015) which initiates mtDNA replication, along with other factors *TFB1M* and *TFB2M*.(Carling et al. 2011; Chiaratti et al. 2011; Grady et al. 2014) Regulators of these transcription factors include *PGC-1α* (peroxisome proliferators-activated receptor gamma coactivator 1 alpha)(Carling et al. 2011; Chiaratti et al. 2011; Harvey et al. 2011) and two nuclear respiratory factors (*NRF-1, NRF-2*).(Carling et al. 2011; Chiaratti et al. 2011; Harvey et al. 2011) Moreover, maintenance of replication requires an adequate mitochondrial nucleotide supply:(González-Vioque et al. 2011) nucleotides may be imported from the cytosol or salvaged by specific mitochondrial enzymes. Defective phosphorylation of deoxyribonucleosides by kinases encoded by *DGUOK* (deoxyguanosine kinase) and *TK2* (thymidine kinase) leads to dysfunctional mitochondrial dNTP synthesis, key regulators of dNTP synthesis in the cytosol include the helicase *C10orf2* (alias *TWINK*),(Bornstein et al. 2008; Rotig and Poulton 2009; Spinazzola et al. 2009; Tyynismaa et al. 2009; Carling et al. 2011; Chiaratti et al. 2011; Harvey et al. 2011; Venegas et al. 2011) along with thymidine phosphorylase (*TYMP*)(Bornstein et al. 2008; Carling et al. 2011; Harvey et al. 2011) and the target of the p53-transcription factor, p53R2 (encoded by *RRM2B*).(Bornstein et al. 2008; Rotig and Poulton 2009; Tyynismaa et al. 2009; Carling et al. 2011; Venegas et al. 2011) The role of succinyl CoA synthase deficiency as a cause of mtDNA depletion is less well understood, but mutations in the alpha and β subunits of Succinyl CoA synthase genes (*SUCLA2, SUCGL1*),(Bornstein et al. 2008; Rotig and Poulton 2009; Spinazzola et al. 2009; Venegas et al. 2011) may be associated with mitochondrial nucleotide depletion.(Rotig and Poulton 2009)

To our knowledge, few genome-wide scans of mtDNA CN have been published, and those that exist are of relatively small sample size,(Curran et al. 2007; López et al. 2014; Workalemahu et al. 2017) or use *in silico* proxies for mtDNA CN without actual biological measurements.(Cai et al. 2015) We had access to directly assayed mtDNA CN in a diverse set of study groups, and so performed hypothesis-generating genome-wide association studies (GWAS) in ~14,000 individuals participants from the Avon Longitudinal Study of Parents and Children (ALSPAC) and the UK Blood Service (UKBS) cohort. For our main analyses, the two most comparable study groups of adult females were combined in a joint analysis (N = 6799, approximately 10-times larger than previous GWASs of directly assayed mtDNA CN),(López et al. 2014; Workalemahu et al. 2017) with results from the other groups presented as opportunistic, secondary analyses. It is known that cellular heterogeneity contributes to mtDNA CN: granulocytes have relatively few mitochondria, whereas lymphocytes are rich in mitochondria, and therefore in mtDNA.(Pyle et al. 2010) Since we also had access to data on white cell proportions, estimated from methylation data in ALSPAC, and assayed directly in UKBS, we performed sensitivity analyses that considered DNA source (whole blood/white cells), and controlled for white cell proportions. Finally, we extracted two SNPs that were robustly related to mtDNA CN in a recent GWAS of mtDNA CN measured *in silico,*(Cai et al. 2015) and compared our results to those published associations.

## Participants and Methods

### Cohort details

ALSPAC is a prospective cohort of mothers and their children. Between 1991-1992, 14,541 women living in the former county of Avon, UK, were recruited during pregnancy, of whom 13,761 were enrolled into the study (women were aged between 16-43 years at recruitment, when samples for mtDNA CN analyses were obtained). Further details are available in the cohort profile papers,(Fraser et al. 2013; Boyd et al. 2013) and the study website contains details of all data that are available through a fully searchable data dictionary: http://www.bris.ac.uk/alspac/researchers/data-access/data-dictionary/. Ethical approval for the study was obtained from the ALSPAC Ethics and Law Committee and the Local Research Ethics Committees.

The UK Blood Service control group are part of the Wellcome Trust Case Control Consortium 2. The UK National Blood Service (UKBS) consists of 3091 unrelated, healthy individuals (aged 17-69 years when samples for mtDNA CN assay were obtained), recruited between September 2005 and February 2006. Informed consent was obtained from all participants in accordance with protocols approved by the Peterborough & Fenland Local Research Ethics Committee in September 2005.

### DNA samples

Blood samples used for mtDNA CN assay were collected from ALSPAC mothers during routine antenatal care. Children included in this study had DNA sampled at either birth (from cord blood, these individuals are hereafter referred to as ALSPAC ‘neonates’), or at a follow-up research clinic assessment at mean age 7 (range: 6-9 years, hereafter ALSPAC ‘6-9 year olds’). Antenatal DNA from mothers was extracted using a phenol-chloroform method.(Jones et al. 2000) DNA from ALSPAC children was extracted using a phenol-chloroform (ALSPAC neonates), or salting-out method (ALSPAC 6-9 year olds).(Jones et al. 2000) DNA sources used for the mtDNA CN assay varied by age group: DNA from whole blood was used for 6-9 year olds, ALSPAC mothers’ DNA was extracted from whole blood or white cells, and ALSPAC neonates had DNA extracted from white cells, as described previously.(Jones et al. 2000)

UKBS blood samples were separated by density centrifugation, and white blood cells were retained to perform DNA extractions, as previously described, using a guanidine-chloroform-based method.(Burton et al. 2007; Nalls et al. 2011) Thus in UKBS, the DNA source was white blood cells for all participants. Blood count information for UKBS samples was provided by Willem Ouwehand at the University of Cambridge as part of an on-going collaboration with Patrick Chinnery. These details are also summarised in **Table 1**.

**Table 1.** Description of study groups and analysis structure in this paper.

| Cohort | Group | % Female | Source of DNA | Age at DNA assay | Site of assay | Covariates in baseline model | Cell proportions source | Cell proportions included in sensitivity model | N (N [CC]) main analysis | N (N [CC]) by subgroups | Meta-analyses (MA) in which each group features |
| --- | --- | --- | --- | --- | --- | --- | --- | --- | --- | --- | --- |
| ALSPAC | Mothers<sup>a</sup>(Antenatal) | 100% | White cells or whole blood [Phenol-Chloroform extracted] | 16-43 years | Bristol | Age, DNA concentration, PC1, PC2, sample type (white cell or whole blood) | Estimated for a subset from Illumina 450k methylation data | Lymphocytes and neutrophils | 5461 (546) (All DNA sources) | 2056 (245) [Whole blood samples]<br>3405 (301) [White cell samples] | MA1 [Main] All mothers<br>MA2 [Sensitivity] White cell mothers only |
|  | Children<sup>b</sup> | 49% | Whole blood [Salt-extracted] | 6-9 years | Bristol | Age, sex, DNA concentration, PC1, PC2 |  | Lymphocytes, monocytes and neutrophils | NA | 2102 (80) | --- |
|  | Neonates<sup>b</sup> | 44% | White cells [Phenol-Chloroform extracted] | 0 years | Bristol | Sex, DNA concentration, PC1, PC2 |  | Lymphocytes, monocytes and granulocytes | NA | 3647 (606) | --- |
| UKBS | Female<sup>a</sup> | 50% | White cells [Guanidine-Chloroform extracted] | 17-69 years | Newcastle | Age, PC1-PC4 | Derived from full blood counts<sup>c</sup> | Lymphocytes and neutrophils | 1338 (1138) [Females only] | --- | MA1 [Main] Females only<br>MA2 [Sensitivity] Females only |
|  | Male |  |  |  |  |  |  |  | NA | 1333 (1122) [Males] | --- |

<sup>a</sup>Denotes group used in main, meta-analyses
<sup>b</sup>Children and neonates in these secondary analyses are unrelated to each other at IBD>0.125, but some are related to the ALSPAC Mothers (see '**Genotype data**' in '**Methods**') <sup>c</sup>See Nalls et al. (2011);(Nalls et al. 2011) Gieger et al. (2011)(Gieger et al. 2011)
CC=cell counts

### Genotype data

#### ALSPAC

ALSPAC Mothers were genotyped on the Illumina Human660W-Quad array (Illumina, San Diego, California, USA) at the Centre Nationale du Génotypage (CNG). ALSPAC children were genotyped with the Illumina HumanHap550-Quad array, by the Wellcome Trust Sanger Institute, Cambridge, UK, and the Laboratory Corporation of America, Burlington, NC, US, using support from 23andMe. Genotypes were called using Illumina GenomeStudio®. Quality control (QC) was performed using PLINK v1.07,(Purcell et al. 2007) phasing using ShapeIT (v2.r644)(O’Connell et al. 2014) and imputation was to the Haplotype Reference Consortium (v1.0), performed using IMPUTE (v3) (http://mathgen.stats.ox.ac.uk/impute/impute.html). The genome build used was GRCh37. Further details of genotype QC are given in **Appendix 1**.

GWAS were run separately in 5,461 ALSPAC mothers, 3,647 6-9 year olds, and 2,102 neonates (see **Appendix 2** for details of selection into the study). Relatedness within each group of participants (mothers, neonates and 6-9 year olds) was assessed by identical-by-descent (IBD) proportions from a genetic-relatedness matrix, calculated using the GCTA standard algorithm,(Yang et al. 2011) based on 1.1 million HapMap3 best-guess tag SNPs, present at a combined allele frequency of >0.01 and imputation quality >0.8 in 17,842 individuals). Within each group (mothers, 6-9 year olds, and neonates) participants were unrelated (IBD>0.125; i.e. first-cousin level). A subset of children were related to the 5,461 ALSPAC mothers: there were 1,611 mother/6-9 year old pairs related at IBD>0.125 (1,570 pairs IBD>0.45) and 869 mother-neonate pairs related at IBD>0.125 (839 IBD>0.45). For some sensitivity analyses, a GWAS of a subset of 2,833 mothers, who are unrelated to any 6-9 year olds or neonates at IBD>0.125 is used. SNPs were filtered by MAF<0.01 and imputation score <0.8 in all study groups,(Marchini and Howie 2010) leaving 7,360,988, 7,410,776 and 7,361,275 SNPs in ALSPAC mothers, 6-9 year olds, and neonates, respectively.

#### UKBS

The UKBS cohort was genotyped using the Illumina 1.2M Duo platform. Raw genotype data (called using Illuminus,(Teo et al. 2007) http://www.sanger.ac.uk/science/tools/illuminus) were downloaded from the European Genotype Archive (http://www.ebi.ac.uk/ega). QC was performed using PLINK v1.90,(Chang et al. 2015) phasing and imputation using EAGLE2 (v2.0.5)(Loh et al. 2016) and PBWT(Durbin 2014) (imputation was to the Haplotype Reference Consortium (v1.0), performed using the Sanger Imputation Server [https://imputation.sanger.ac.uk/]). The genome build was GRCh37. Further details are given in **Appendix 1**.

GWAS were run in 2,671 UKBS individuals (1,333 males, 1,338 females). Individuals were unrelated at any level of IBD. After filtering by MAF<0.01 and imputation quality >0.8, 7,441,490 variants remained (7,369,986 males only, 7,373,492 females only).

#### Assay of mtDNA CN

For mtDNA CN assay details in ALSPAC and UKBS, see **Appendix 2.** Both cohorts had mtDNA CN assayed by quantitative PCR. Both assays used *B2M* as the single-gene reference, but mtDNA amplicons differed. Raw data are plotted in **Supplementary Figure 1**.

Despite differences in raw mtDNA CNs between study groups, validation analyses of the two adult cohorts suggested relative mtDNA CNs were reliable: Cross-validation of qPCR methodology between centres was performed on 384 random samples (169 from ALSPAC, 185 from UKBS). Samples were assayed by PG and AG in Bristol, and RB in Newcastle, using cohort-specific protocols. There was moderate-to-good agreement between z-scores of mtDNA CNs assayed in ALSPAC by AG and PG (r[Spearman] = 0.68), and between those obtained from the two sets of exchanged plates (ALSPAC r[Spearman] = 0.58 [analysts: AG/RB] and UKBS r[Spearman] = 0.69 [analysts: PG/RB]) (**Supplementary Figure 2**). However, Panel B of **Supplementary Figure 2** suggests that there may have been some non-linearity when comparing the two assays. To control for absolute differences in mtDNA CNs, z-scored phenotypes were used in GWAS (after log-transformation to approximate normality). Z-scores were computed separately for ALSPAC mothers, 6-9 year olds, neonates, and UKBS.

#### Statistical analysis

##### Genome-wide association study

GWAS were undertaken separately for ALSPAC mothers, ALSPAC 6-9 year olds, ALSPAC neonates, UKBS females, and UKBS males. Additive models were fitted, using dosage data for ALSPAC, and best-guess data for UKBS. The genome-wide significance threshold was *p*=5e-08.(Aschebrook-Kilfoy et al. 2015)

###### Main analyses

We had access to several study groups with relevant data. Since these groups were diverse in nature, and were of relatively small sample sizes (compared to some complex trait GWAS), we did not consider them as ‘discovery’ and ‘replication’ cohorts. Instead, after validating and harmonising data (see ‘**Assay of mtDNA CN**’), we considered our main analyses to be hypothesis-generating GWAS of the two most comparable groups, i.e. all adult females (5461 ALSPAC mothers and 1338 UKBS females). This decision also took into account results from some preliminary analyses that suggested some possible differences between UKBS females and males (see **Supplementary Figures 3 and 4**), although we acknowledge that we have insufficient power to detect sex differences that are not potentially due to chance.

Thus, results from ALSPAC mothers and UKBS females were meta-analysed, using random-effects models (‘Meta-analysis 1’). Since ALSPAC mothers had DNA extracted from two sources, a sensitivity meta-analysis (‘Meta-analysis 2’, N=4,743) restricted the ALSPAC mothers to 3,405 females with white cell DNA extracted by a phenol-chloroform method (i.e. the most comparable subgroup to UKBS females, all of whom had white cell DNA extracted by a guanidine-chloroform based method(Burton et al. 2007; Nalls et al. 2011)). Heterogeneity was assessed using Cochran’s Q statistic,(Cochran 1954) and the I^2^ statistic.(Higgins et al. 2003)

###### Secondary analyses

Results from GWAS of 3,647 ALSPAC children [6-9 years], 2,102 ALSPAC neonates, and 1333 UKBS males are presented as secondary analyses. We applied the same p-value threshold for genome-wide significance, and also specifically looked at whether hits identified in these groups showed similar directions and magnitudes of association in the meta-analyses of adult females. However, we did not consider these groups to be suitable replication samples for the main analyses of adult females, given their small sample sizes, and sex and age differences.

###### Look-up of top loci from a recent large GWAS of in silico mtDNA CN

A recent large GWAS (using a discovery sample of 10,560 Han Chinese females) identified *TFAM* (mitochondrial transcription factor A) and *CDK6* (Cyclin Dependent Kinase 6), as loci strongly associated with mtDNA CN estimated *in silico*, from sequence data.(Cai et al. 2015) The replication sample for that study included 1,753 ALSPAC children within the UK10K consortium.(Walter et al. 2015) There is overlap between that group, and the 6-9 year olds studied in the current analysis.(Cai et al. 2015) Using all available unrelated participants in the current study (i.e. excluding mother-child duos, N=11,253 total), we meta-analysed the two lead SNPs at these loci, and compared our results to those of this previous GWAS.(Cai et al. 2015)

###### Covariates

Covariates included age at mtDNA CN assay, sex, and DNA concentration (ng/μL)(Malik et al. 2011) (ALSPAC only, measured by a PicoGreen^®^ fluorescence-based method), and principal components (2 in ALSPAC, 4 in UKBS), to adjust for population structure. The ALSPAC mothers’ analysis was adjusted for DNA source (white cells or whole blood) (see **Error! Reference source not found.**).

For all analyses, effect sizes adjusted for cell-proportions are also presented, since cell lineages vary in their average numbers of mitochondria.(Pyle et al. 2010) In ALSPAC, cell proportions have been estimated in the ‘Accessible Resource for Integrated Epigenomics’ (ARIES) subset(Relton et al. 2015), using DNA methylation data from the Illumina Infinium HumanMethylation450 BeadChip (450K) array, and the Houseman method.(Houseman et al. 2012) 546 mothers, 606 6-9 year olds, and 82 neonates included in this study had cell proportion data. Proportions were estimated from methylation data derived from the same time points as mtDNA CNs were assayed (antenatally for mothers, at birth and ~7 years for children). In this study, we only performed cell proportion adjusted analyses if participants had mtDNA CN assayed in the same DNA sample type that was used for cellular proportion estimation (i.e. white cells/whole blood).(Faul et al. 2007) Lymphocytes (total of CD8T, CD4T, B lymphocytes) and neutrophils (granulocytes for neonates(Cardenas et al. 2016)) were included as covariates for mothers, with the addition of monocytes for 6-9 year olds and neonates (monocytes were not used in the mothers’ analyses since this prevented model convergence). In UKBS, neutrophil and lymphocyte proportions derived from full blood count data were included.(Nalls et al. 2011; Gieger et al. 2011)

###### Software

GWAS were performed with SNPTESTv2.5.0,(Marchini et al. 2007) and meta-analysed with META v1.7.0.(Liu et al. 2010b) The ‘qqman’ R package [http://cran.r-project.org/web/packages/qqman/] was used to create Manhattan/quantile-quantile (QQ) plots. ANNOVAR(Wang et al. 2010) was used to annotate loci [http://annovar.openbioinformatics.org/en/latest/], and the ‘bedtools’ ‘clusterBed’ function [https://github.com/arq5x/bedtools2](Quinlan and Hall 2010) was used to cluster loci. Regional association plots were produced with LocusZoom v1.3, using an hg19 reference (1000G March 2012).(Pruim et al. 2010)

###### Power

Genetic Power Calculator(Sham et al. 2000; Purcell et al. 2003) was used to determine the minimum detectable effect sizes at 80% power, given the sample size of the study groups used in the main meta-analyses (“http://www.cureffi.org/2012/12/05/power-for-gwas-and-extreme-phenotype-studies/”). This method requires ‘Total QTL variance’ (i.e. the proportion of variance in a quantitative trait locus, *V_q_* explained by the causal variant) as input. Assuming a standardised normal distribution (phenotype in SD units), the following is true:

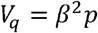

where *β* is the effect size (in SD units), and *p* is the minor allele frequency (MAF). Estimated minimal effect sizes for our given sample sizes, for a range of *V_q_* values were calculated, with a minimum value of 0.001 (equivalent to an effect size of 0.316 at MAF=0.01, and 0.045 at MAF=0.5). Linkage disequilibrium (LD) of 0.8 (measured by the D’ metric) was assumed between causal and tag variants. Power curves are shown in **Supplementary Figure 5**. Heat maps of power by effect size and MAF are shown in **Supplementary Figure 6**. For the largest (N=6799 [5461+1338]) and smallest (N=1333) GWAS undertaken in this paper (i.e. the main meta-analysis of adult females, and the smallest secondary analysis of UKBS males, respectively), minimum *V_q_* values detectable were 0.0091 and 0.0458. At MAF=0.25, this is equivalent to effect sizes of 0.191 and 0.428. However, it should be noted that these calculations do not take account of measurement error in mtDNA CN, white cell heterogeneity, covariate adjustment, or the use of random-effects meta-analysis for the main GWAS.

## Results

### Main analysis

Manhattan/QQ plots for GWAS of ALSPAC mothers (for all 5,461 mothers, and for 3,405 with white cell-extracted DNA), UKBS females, and the two meta-analyses are shown in **Figure 1** and **Figure 2**. Regional association plots for loci identified from the meta-analyses are in **Figure 3**. For strongest associations from separate GWAS of ALSPAC Mothers and UKBS Females, see **Supplementary Tables 1-5** and **Appendix 3**).

**Figure 1.**
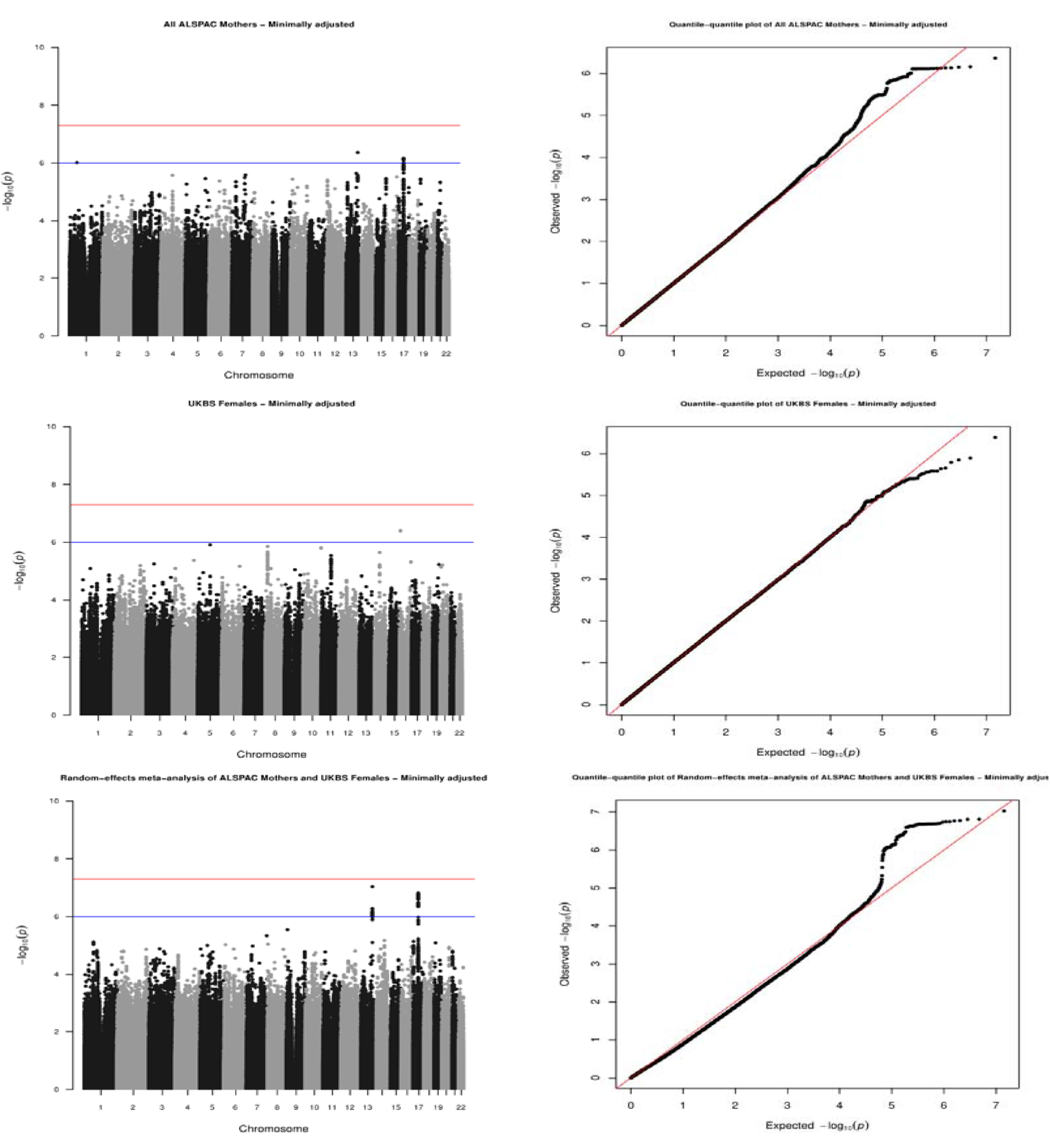
Manhattan (left) / QQ plots (right) for ALSPAC (all Mothers) and UKBS (females), and random-effects meta-analysis of both cohorts. Top row=ALSPAC, Middle row=UKBS Females, Bottom row=Meta-analysis (random-effects). λ=0.995 and 1.011 for ALSPAC (all Mothers) and UKBS Females respectively, and meta-analyses are corrected for these lambdas. ‘Minimally-adjusted’ refers to the fact that these results are from the analysis that did not adjust for cell proportions.

**Figure 2.**
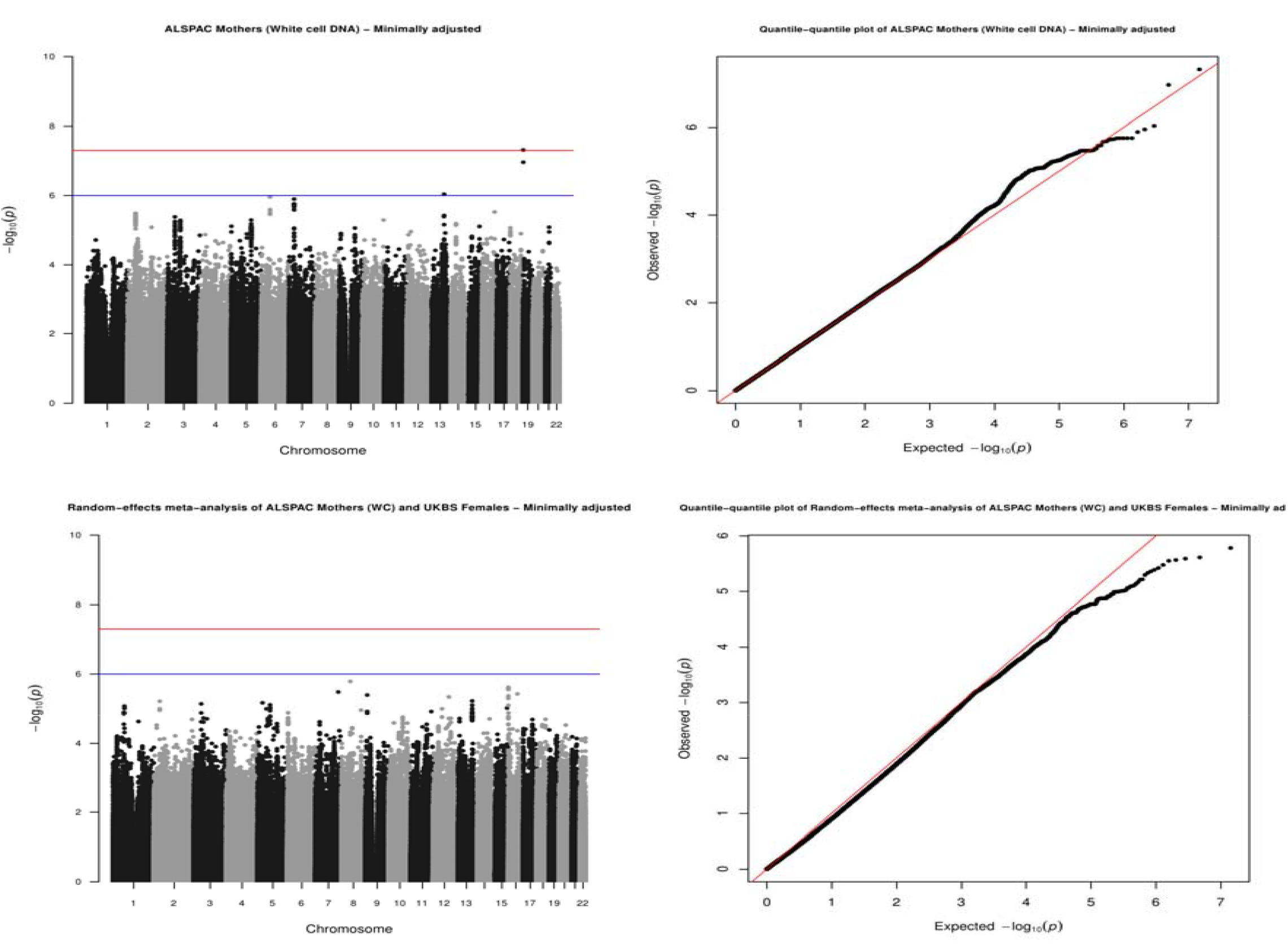
Manhattan (left) / QQ plots (right) for ALSPAC (white cell Mothers) and random-effects meta-analysis of both cohorts. Top row=ALSPAC, Bottom row=Meta-analysis (random-effects). λ=0.992 and 1.011 for ALSPAC (white cell Mothers) and UKBS females (see other plot), and meta-analyses are corrected for these lambdas. ‘Minimally-adjusted’ refers to the fact that these results are from the analysis that did not adjust for cell proportions.

**Figure 3.**
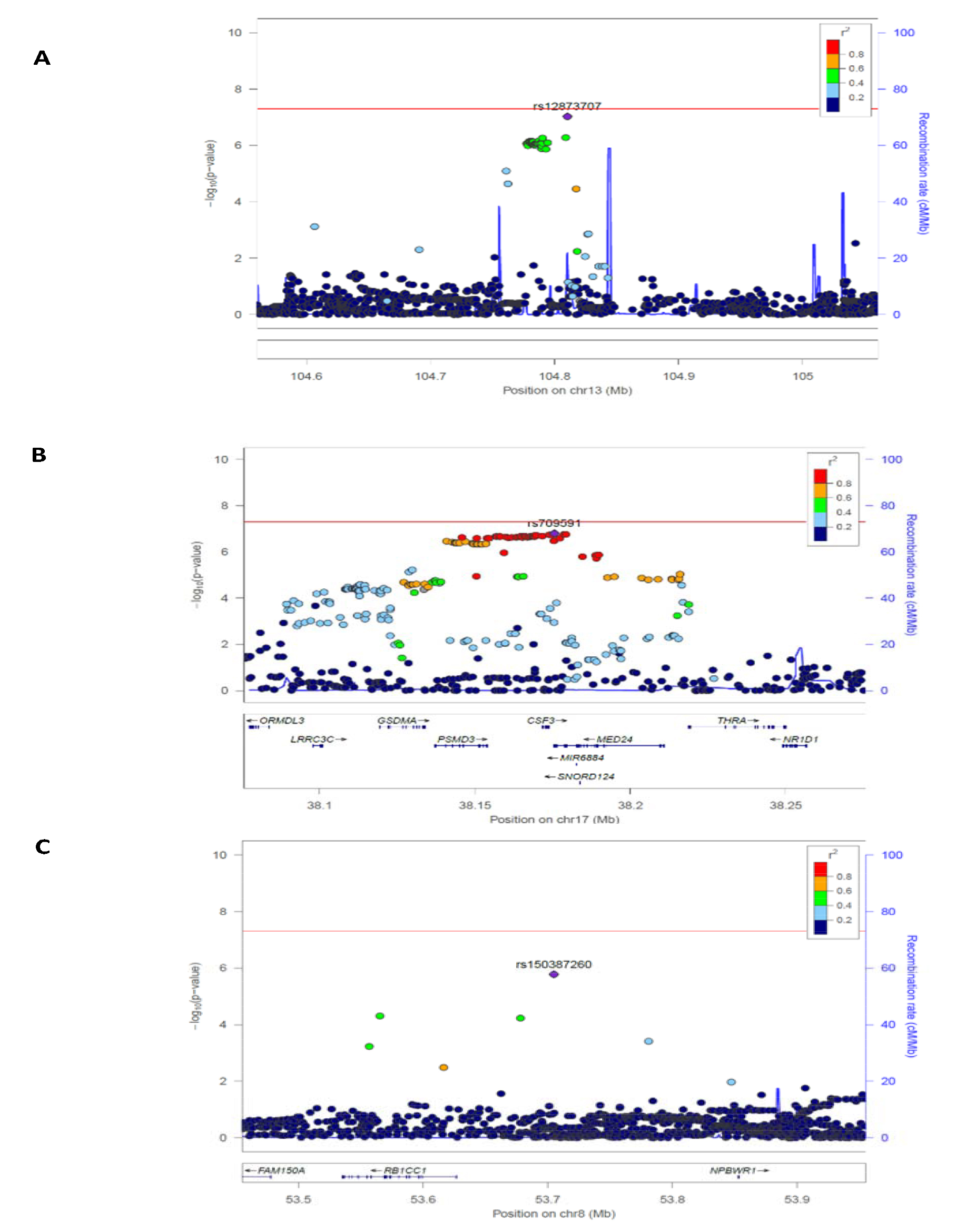
Regional association plots (created with LocusZoom) of loci presented in the meta-analyses (**Table 2**). Panels A-B are of loci identified in the meta-analysis of all ALSPAC mothers and UKBS females; panels C is of the one locus identified after restriction of the meta-analysis to ALSPAC mothers with DNA extracted from white cells, only. The lead SNP is annotated in purple, with other SNPs colour coded according to values of LD. P-value and recombination level are shown on the left and right y axes. A schematic of the region, along with coordinates and annotations (if any) is shown at the bottom of each plot. See also **Table 2**.

Results from the main meta-analysis of adult females (N=6799, ‘Meta-analysis 1’), as well as the analysis restricted to those with white cell-extracted DNA (N=4743, ‘Meta-analysis 2’) are given in **Table 2**. SNPs in associated regions were clustered into 1Mb windows.(Quinlan and Hall 2010) For each analysis, results are shown before and after cell-proportion adjustment. Annotations are given for loci within 200kb.(Eberle et al. 2006) Coordinates are hg19.

**Table 2.**
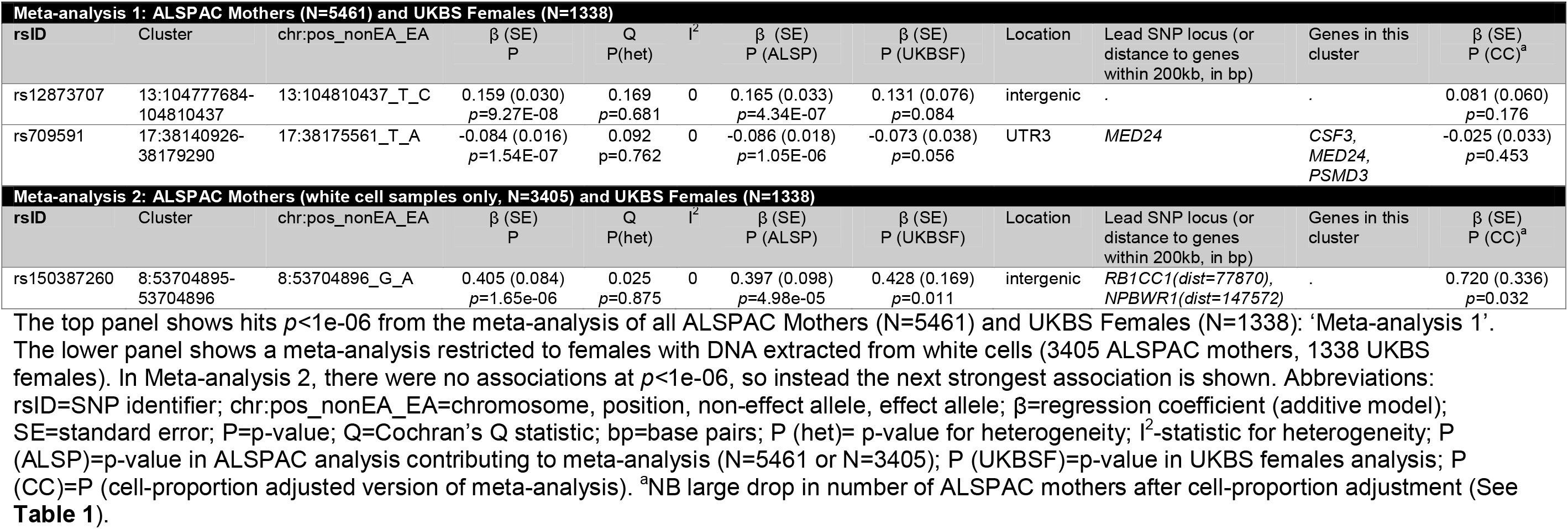
Summary statistics for top SNPs associated with mtDNA CN in two random-effects meta-analyses.

| **Meta-analysis 1: ALSPAC Mothers (N=5461) and UKBS Females (N=1338)** |  |  |  |  |  |  |  |  |  |  |  |
| --- | --- | --- | --- | --- | --- | --- | --- | --- | --- | --- | --- |
| rsID | Cluster | chr:pos_nonEA_EA | $\beta$ (SE)<br>P | Q<br>P(het) | $I^2$ | $\beta$ (SE)<br>P (ALSP) | $\beta$ (SE)<br>P (UKBSF) | Location | Lead SNP locus (or<br>distance to genes<br>within 200kb, in bp) | Genes in this<br>cluster | $\beta$ (SE)<br>P (CC)<sup>a</sup> |
| rs12873707 | 13:104777684-104810437 | 13:104810437_T_C | 0.159 (0.030)<br>$p=9.27E-08$ | 0.169<br>$p=0.681$ | 0 | 0.165 (0.033)<br>$p=4.34E-07$ | 0.131 (0.076)<br>$p=0.084$ | intergenic | . | . | 0.081 (0.060)<br>$p=0.176$ |
| rs709591 | 17:38140926-38179290 | 17:38175561_T_A | -0.084 (0.016)<br>$p=1.54E-07$ | 0.092<br>$p=0.762$ | 0 | -0.086 (0.018)<br>$p=1.05E-06$ | -0.073 (0.038)<br>$p=0.056$ | UTR3 | MED24 | CSF3,<br>MED24,<br>PSMD3 | -0.025 (0.033)<br>$p=0.453$ |
| **Meta-analysis 2: ALSPAC Mothers (white cell samples only, N=3405) and UKBS Females (N=1338)** |  |  |  |  |  |  |  |  |  |  |  |
| rsID | Cluster | chr:pos_nonEA_EA | $\beta$ (SE)<br>P | Q<br>P(het) | $I^2$ | $\beta$ (SE)<br>P (ALSP) | $\beta$ (SE)<br>P (UKBSF) | Location | Lead SNP locus (or<br>distance to genes<br>within 200kb, in bp) | Genes in this<br>cluster | $\beta$ (SE)<br>P (CC)<sup>a</sup> |
| rs150387260 | 8:53704895-53704896 | 8:53704896_G_A | 0.405 (0.084)<br>$p=1.65e-06$ | 0.025<br>$p=0.875$ | 0 | 0.397 (0.098)<br>$p=4.98e-05$ | 0.428 (0.169)<br>$p=0.011$ | intergenic | RB1CC1(dist=77870),<br>NPBWR1(dist=147572) | . | 0.720 (0.336)<br>$p=0.032$ |

The top panel shows hits $p < 1e-06$ from the meta-analysis of all ALSPAC Mothers (N=5461) and UKBS Females (N=1338): 'Meta-analysis 1'.
The lower panel shows a meta-analysis restricted to females with DNA extracted from white cells (3405 ALSPAC mothers, 1338 UKBS females).
In Meta-analysis 2, there were no associations at $p < 1e-06$ , so instead the next strongest association is shown. Abbreviations:
rsID=SNP identifier; chr:pos\_nonEA\_EA=chromosome, position, non-effect allele, effect allele; $\beta$ =regression coefficient (additive model);
SE=standard error; P=p-value; Q=Cochran's Q statistic; bp=base pairs; P (het)= p-value for heterogeneity; $I^2$ -statistic for heterogeneity; P
(ALSP)=p-value in ALSPAC analysis contributing to meta-analysis (N=5461 or N=3405); P (UKBSF)=p-value in UKBS females analysis; P
(CC)=P (cell-proportion adjusted version of meta-analysis). <sup>a</sup>NB large drop in number of ALSPAC mothers after cell-proportion adjustment (See
Table 1).

#### Meta-analysis 1: all adult females (N=6799)

No SNP passed the genome-wide significance threshold of *p*<5e^−08^. The top panel of **Table 2** describes two top loci (*p*<1e-06) identified from the main meta-analysis of all adult females: these included an intergenic locus on chr13 (lead SNP rs12873707, β [SE] 0.159 [0.030], *p*=9.27e-08, I^2^=0), and a locus on chr17 (lead SNP rs709591, β [SE] −0.084 (0.016) *p*=1.54e-07, I^2^=0). A list of all SNPs associated at *p*<1e-06 in these loci is given in **Supplementary Table 6**.

Regional association plots (see Panels A and B of **Figure 3**) show the LD structure at these loci. Panel A shows that there are no nearby SNPs in LD with the lead chr13 variant. Panel B shows a large region of LD at the chr17 locus, spanning the genes *PSMD3, CSF3,* and *MED24*. SNPs in this region are associated with neutrophil count:(Nalls et al. 2011) given this fact, and since cellular heterogeneity affects mtDNA CN,(Pyle et al. 2010; Xu et al. 2013; Tin et al. 2016) this locus was assessed in more detail, by extracting the SNP from all study groups (**Table 3**). Despite differences in the proportion of participants with information that enabled adjustment for cell-type, there was consistency across study groups with each showing an approximate 50% attenuation of the effect size with adjustment for cell proportions.

**Table 3.** General concordance of association at rs709591, before adjustment for cell counts

| Cohort | Group | Subgroup | P | Info score<br>(Imputation<br>quality) | $\beta$ | SE | N |
| --- | --- | --- | --- | --- | --- | --- | --- |
| **Unadjusted** |  |  |  |  |  |  |  |
| **ALSPAC<sup>a</sup>** | Mothers | All | 1.05E-06 | 0.998 | -0.086 | 0.018 | 5461 |
|  |  | White cell samples | 6.26E-04 | 0.998 | -0.081 | 0.024 | 3405 |
|  | Children<sup>b</sup> | All | 0.524 | 0.998 | -0.015 | 0.024 | 3647 |
|  | Neonates<sup>b</sup> | All | 1.36E-04 | 0.998 | -0.119 | 0.031 | 2102 |
| **UKBS** | All | Females | 0.056 | 0.999 | -0.073 | 0.038 | 1338 |
|  |  | Males | 0.025 | 0.999 | -0.090 | 0.040 | 1333 |
| **Cell-proportion adjusted (see Table 1 for covariates)** |  |  |  |  |  |  |  |
| **ALSPAC<sup>a</sup>** | Mothers | All | 0.829 | 0.997 | 0.012 | 0.057 | 546 |
|  |  | White cell samples | 0.774 | 0.999 | 0.024 | 0.085 | 301 |
|  | Children<sup>b</sup> | All | 0.702 | 0.998 | 0.019 | 0.051 | 606 |
|  | Neonates<sup>b</sup> | All | 0.661 | 1.000 | -0.08 | 0.18 | 82 |
| **UKBS** | All | Females | 0.278 | 0.999 | -0.044 | 0.041 | 1138 |
|  |  | Males | 0.241 | 0.999 | -0.049 | 0.042 | 1122 |

<sup>a</sup>NB large drop in power for ALSPAC (See **Table 1**).
<sup>b</sup>Some children in these analyses are related to ALSPAC Mothers, however, in N=2833 unrelated ALSPAC mothers, this association persisted ( $\beta$ [SE] -0.120 [0.024], $p=4.01\text{e-}07$ ).

#### Meta-analysis 2: adult females with white cell-extracted DNA (N=4743)

The bottom panel of **Table 2** gives details of the strongest association in the meta-analysis restricted to ALSPAC mothers with DNA extracted from white cells (‘Meta-analysis 2’). The locus associated with mtDNA CN (*p*<1e^−06^) was rs150387260, an intergenic variant (β [SE] 0.405 [0.084], *p*=1.65e-06, I^2^=0, see also **Supplementary Table 7**). A regional association plot (panel C of **Figure 3**) shows that few neighbouring SNPs are in substantial LD with this SNP. The nearest gene was RB1 Inducible Coiled-Coil 1 (*RB1CC1*), 78kb upstream. The product of this tumour suppressor gene is implicated in cell growth, migration, proliferation, apoptosis, and autophagy.(Stelzer et al. 2016) *RB1CC1* deletions are associated with increased numbers of mitochondria in hematopoietic stem cells(Liu et al. 2010a) and mice,(Yao et al. 2015) and with breast cancer in humans.(Chano et al. 2002)

#### Effect of adjusting for cell-proportions in these two meta-analyses

In Meta-analysis 1 (all adult females, regardless of DNA source), point estimates attenuated after cell proportion adjustment, although the reduced sample size meant that confidence intervals were wide. In contrast, after cell proportion adjustment for the association identified from Meta-analysis 2 (restricted to females with white cell-extracted DNA), confidence intervals were also wide, but the effect estimate was not attenuated (it increased slightly).

### Secondary analyses

**Table 4** shows results of GWAS in ALSPAC 6-9 year olds, neonates, and UKBS males. Only genome-wide significant loci are shown (for UKBS males, top loci at *p*<1e-06 are shown, as there were no loci *p*<5e-08. When comparing effect sizes between any hits in these groups with effects of the same SNPs in the main (adult female) analyses, it should be noted that there are ~1600 mother-child duos between ALSPAC mothers included in the main analysis and the 6-9 year olds / neonates (see ‘**Methods**’). Results of the associations of SNPs at a known neutrophil count locus (*PSMD3, CSF3, MED24*) identified in our main analyses in these secondary analysis groups are discussed above (see ‘Main analysis’).

**Table 4.** Lead SNPs in secondary analyses

| ALSPAC 6-9 year olds (N=3647) |  |  |  |  |  |  |  |  |  |  |
| --- | --- | --- | --- | --- | --- | --- | --- | --- | --- | --- |
| rsID | Cluster | chr:pos_nonEA_EA | $\beta$ (SE)<br>P | $\beta$ (SE)<br>P (Proxy) | Location | Nearby<br>genes | Lead SNP locus (or<br>distance to genes<br>within 200kb, in bp) | $\beta$ (SE)<br>P (CC)<sup>a</sup> | $\beta$ (SE)<br>P (MA1) | $\beta$ (SE)<br>P (MA2) |
| rs139045492 | 20:22836201-22836202 | 20:22836202_G_A | -0.695 (0.119)<br>$p=6.35E-09$ | -- | intergenic | . | *SSTR4*<br>(dist=179855) | -0.489 (0.240)<br>$p=0.042$ | 0.080 (0.078)<br>$p=0.306$ | -0.005 (0.097)<br>$p=0.956$ |
| rs144874692 | 3:175689046-175689047 | 3:175689047_G_A | -0.632 (0.112)<br>$p=1.78E-08$ | -0.328 (0.105)<br>$p=0.002$ | intergenic | . | *NAALADL2*<br>(dist=165619) | -0.024 (0.263)<br>$p=0.926$ | Proxy:<br>0.00 (0.075)<br>$p=1.00$ | Proxy:<br>0.016 (0.094)<br>$p=0.861$ |
| ALSPAC Neonates (N=2102) |  |  |  |  |  |  |  |  |  |  |
| rsID | Cluster | chr:pos_nonEA_EA | $\beta$ (SE)<br>P | $\beta$ (SE)<br>P (Proxy) | Location | Nearby<br>genes | Lead SNP locus (or<br>distance to genes<br>within 200kb, in bp) | $\beta$ (SE)<br>P (CC)<sup>a</sup> | $\beta$ (SE)<br>P (MA1) | $\beta$ (SE)<br>P (MA2) |
| rs10424198 | 19:17354824-17461934 | 19:17409671_C_T | -0.262 (0.034)<br>$p=1.40E-14$ | -- | intronic | *ABHD8*,<br>*ANKLE1*,<br>*ANO8*,<br>*BABAM1*,<br>*DDA1*,<br>*GTPBP3*,<br>*MRPL34*,<br>*NR2F6*,<br>*PLVAP*,<br>*USHBP1* | *ABHD8* | -0.326 (0.193)<br>$p=0.095$ | -0.010 (0.017)<br>$p=0.556$ | 0.002 (0.022)<br>$p=0.911$ |
| UKBS (Males) (N=1333) |  |  |  |  |  |  |  |  |  |  |
| rsID | Cluster | chr:pos_nonEA_EA | $\beta$ (SE)<br>P | $\beta$ (SE)<br>P (Proxy) | Location | Nearby<br>genes | Lead SNP locus (or<br>distance to genes<br>within 200kb, in bp) | $\beta$ (SE)<br>P (CC) | $\beta$ (SE)<br>P (MA1) | $\beta$ (SE)<br>P (MA2) |
| rs11687359 | 2:1969703-1969704 | 2:1969704_G_A | 0.401 (0.078)<br>$p=2.94e-07$ | -- | intronic | *MYT1L* | *MYT1L* | 0.362 (0.080)<br>$p=6.52e-06$ | 0.006 (0.032)<br>$p=0.845$ | 0.043 (0.041)<br>$p=0.286$ |
| rs13070764 | 3:73309902-73310337 | 3:73309903_G_A | -0.497 (0.099)<br>$p=5.84e-07$ | -- | intergenic | . | *PPP4R2*<br>(dist=194892),<br>*PDZRN3*<br>(dist=121678) | -0.509 (0.100)<br>$p=4.15e-07$ | 0.100 (0.043)<br>$p=0.020$ | 0.098 (0.060)<br>$p=0.101$ |

Abbreviations: rsID=SNP identifier; chr:pos\_nonEA\_EA=chromosome, position, non-effect allele, effect allele; bp=base pairs; $\beta$ =regression coefficient (additive model); SE=standard error; P=p-value; P (Proxy)=p-value (for a proxy SNP, rs144928561, $r^2=0.29$ ); P (CC)=p-value (cell-proportion adjusted analysis); P (MA1)=p-value ('Meta-analysis 1', 6799 adult females); p-value (MA2)=P ('Meta-analysis 2', 4743 females with DNA from white cells). <sup>a</sup>NB large drop in power for ALSPAC mothers (See **Table 1**).

#### Associations in ALSPAC 6-9 year olds

There were two genome-wide significant, intergenic loci in 6-9 year olds (rs139045492 on chr20, β [SE] −0.695 [0.119], *p*=6.35e-09 and rs144874692 on chr3, β [SE] −0.632 [0.112] *p*=1.78e-08). The rs139045492 association persisted after cell proportion adjustment. rs139045492 was not directionally concordantly (or statistically significantly) associated with mtDNA CN in either of the adult female main meta-analyses. The other association (rs144874692), which attenuated after cell-proportion adjustment, was not available in the main analysis. Therefore, a proxy (rs144928561, r^2^=0.29, β [SE] −0.328 [0.105], *p*=0.002 in ALSPAC 6-9 year olds) was used to explore whether there was any evidence of it being associated with mtDNA CN in the main analyses. This proxy SNP was not directionally concordantly associated in either main meta-analysis. Manhattan/QQ plots are available in **Supplementary Figure 7**. A list of all SNPs at *p*<1e-06 in the 6-9 year olds is available in **Supplementary Table 8**.

#### Associations in ALSPAC neonates

The locus most strongly associated with mtDNA CN in the ALSPAC neonates was at chromosome 19, in *ABHD8* (abhydrolase-domain containing 8) (lead SNP: rs10424198, β [SE] −0.262 (0.034), *p*=1.40e-14). Sixteen SNPs were identified at genome-wide significance within *BABAM1, ANKLE1, ABHD8* and *MRPL34*. Whilst the standard error for this SNP was large after cell-count adjustment, this latter analysis was in a tiny sample (N=82), and yet still the point estimate remained broadly consistent (cell-proportion adjusted β [SE] −0.326 [0.193], *p*=0.095). However, there was no evidence of a concordant association in either meta-analysis from the main analyses. Manhattan/QQ plots are available in **Supplementary Figure 8**. A list of SNPs associated with mtDNA CN at *p*<1e-06 in neonates is available in **Supplementary Table 9**.

#### Associations in UKBS (males)

No SNPs were associated at *p*<5e-08 in UKBS males. Two SNPs were associated at *p*<1e-06: an intronic SNP in *MYT1L* (Myelin Transcription Factor 1 Like, β (SE) 0.401 (0.078), *p*=2.94e-07), and two intergenic SNPs 121kb upstream of *PDZRN3* (PDZ Domain Containing Ring Finger 3, lead SNP β [SE] −0.497 [0.099], *p*=5.84e-07). Whilst these SNPs survived cell proportion adjustment, neither locus showed evidence of an association concordant in terms of direction and magnitude in either main meta-analysis. Manhattan and QQ plots for this analysis are available in **Supplementary Figure 3**. A list of SNPs at *p*<1e-06 in UKBS males is available in **Supplementary Table 10**.

### Look-up of top loci from a recent large GWAS of in silico mtDNA CN

The results of a look-up of two loci (rs11006126 and rs445) that are strongly associated with mtDNA CN measured *in silico*(Cai et al. 2015) are presented in **Figure 4**. Estimates for these SNPs were extracted from, and meta-analysed across all study groups (after removing ALSPAC mother-offspring pairs; total N for these meta-analyses = 11253, see also ‘**Genotype data’**). rs11006126 was associated with mtDNA CN across our study groups (β [SE] 0.046 [0.017], *p*=0.007, but the effect size was considerably (~1/7 of the size) smaller than that observed in the discovery and replication cohorts of the *in silico* GWAS. There was no evidence of replication of rs445 across our study groups (β [SE] 0.021 [0.022], *p*=0.328) compared with discovery and replication result from the *in silico* GWAS.

**Figure 4.**
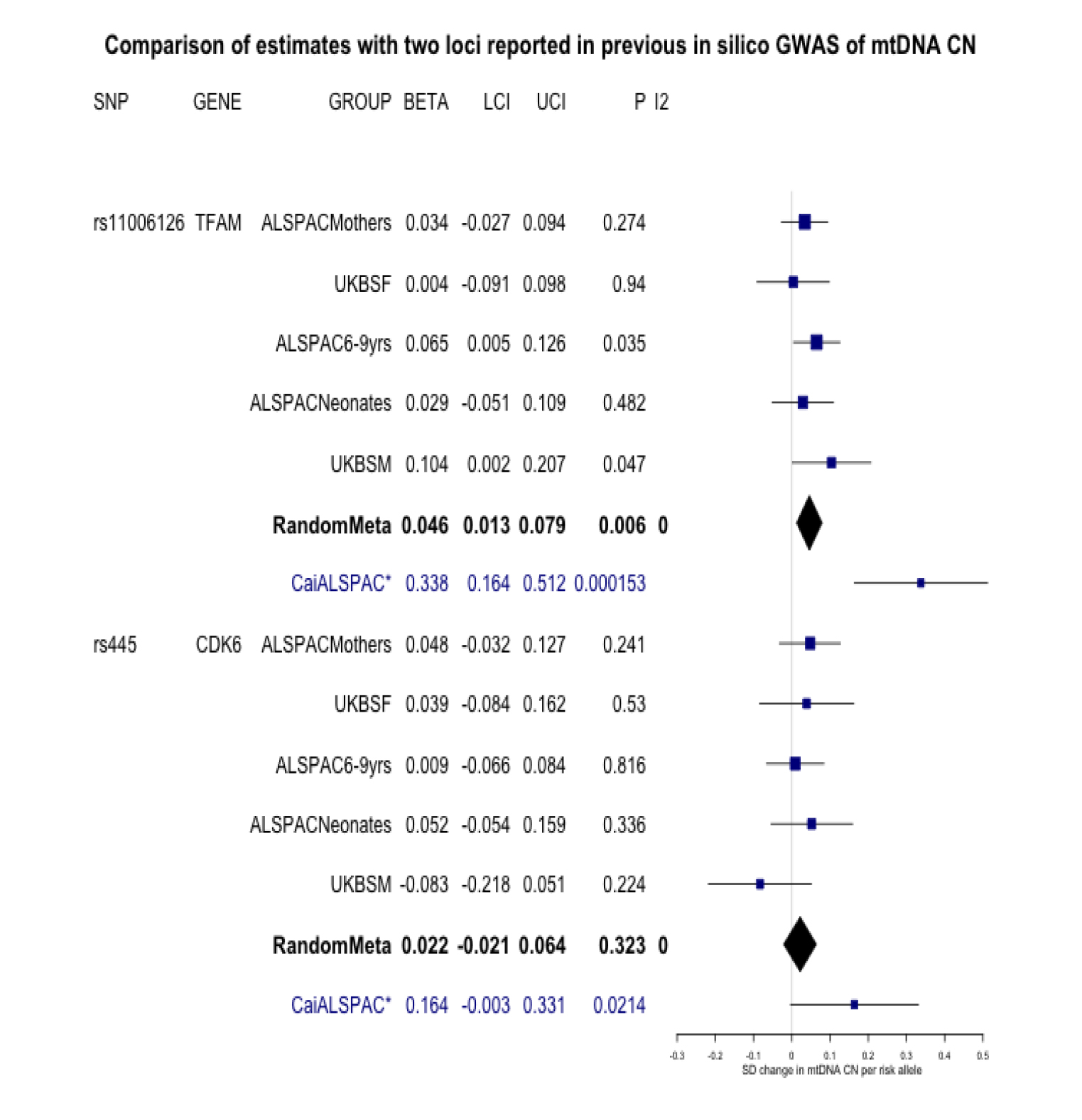
Extraction and meta-analysis of two top loci from Cai *et al.* (2015) from study groups in this GWAS. SNPs for two loci identified in a GWAS of *in silico* estimated mtDNA CN were extracted from each of the study groups in this cohort (NB: a smaller subset of 2833 ALSPAC mothers were used, since there were mother-child duos present between the original study group of 5461 and the two groups of ALSPAC children) Total N=11253. Columns: SNP=rsID; Gene=gene name; Group=study group, Beta=effect size; LCI/UCI=95% confidence interval (lower, then upper bound), P=p value, and I2=I^2^ metric for heterogeneity. Meta-analyses were by random-effects, and are shown as black diamonds. For reference, the ALSPAC estimate from Cai *et al.* (2015) is shown for each locus. This replication group included ALSPAC 6-9 year olds (with mtDNA CN assayed from sequence data). Betas had to be harmonised, as those inCai *et al.* [2015] were given as SD change in mtDNA CN per SD increase in genotype. SD of genotype was estimated from allele frequencies provided for the cohort byCai *et al.* [2015] given as 0.342 for rs445 and 0.169 for rs11006126 (in the supplement of this paper). SDs were then calculated as sqrt(2*(1-MAF)*MAF) (evaluating to 0.53 and 0.67 for rs11006126 and rs445, respectively). Betas and standard errors were then transformed from those given in Table 1 of Cai *et al.* to a ‘per risk allele’ scale, by multiplying the given beta by (1/the estimated SD) i.e. rs11006126=0.179*(1/0.53); rs445=0.110*(1/0.67). CaiALSPAC=Result from Cai et al. (2015). UKBSF/UKBSM=Females and Males in UKBS cohort.

## Discussion

We conducted the largest ever GWAS of directly assayed mtDNA CN, in two population-based cohorts. Although diversity between our study groups prevented us from conceptualising a traditional discovery and replication model, we considered our study as hypothesis generating, and meta-analysed the two most comparable groups (ALSPAC mothers and UKBS females) as our main analyses, and other subgroups as opportunistic, secondary analyses.

There were no genome-wide significant hits in the meta-analysis of adult females, but SNPs at a known neutrophil count locus (*PSMD3, CSF3, MED24*)(Okada et al. 2010) were associated with mtDNA CN at p~2e-07. It is established that cellular heterogeneity is related to mtDNA CN,(Pyle et al. 2010) and one of the strengths of this study is that we were able to control for this. This chr17 locus showed notable consistency of effect sizes across several study groups (UKBS males, UKBS females, ALSPAC neonates), corresponding to a ~0.08 reduction in SD units of log mtDNA CN per risk allele. However, there was notable (~50%) attenuation with adjustment for white cell proportions. Thus, a key finding of this study is the importance of undertaking GWAS of directly assayed mtDNA CN in sample sizes that are much larger, and that also have measures of white cell proportions. Ideally, these studies should be sufficiently large to enable exploration of any possible variation in nuclear genetic control of mtDNA copy number variation by sex and age.

Secondary analyses in ALSPAC children (6-9 years) and ALSPAC neonates, and UKBS males revealed three genome-wide significant hits: two intergenic loci in ALSPAC 6-9 year olds, and a region containing *ABDH8* (abhydrolase-domain containing-8, a gene in head-to-head orientation with mitochondrial ribosomal protein L34, *MRPL34*) in ALSPAC neonates. None of these associations were at all evident in the main meta-analysis of adult females. Another mitochondrial ribosomal protein, *MRPL37* has previously been found to associate with mtDNA CN, which (López et al. 2014) raises the possibility that the *BABAM1-ANKLE1-ABHD8-MRPL34* locus might be a true signal, despite the lack of evidence of effect in our main analyses. Mitochondrial ribosomal proteins are encoded by nuclear DNA, synthesized on cytoplasmic ribosomes, then imported into mitochondria, where they facilitate the translation of mitochondrial mRNA, in conjunction with the two mitochondrially encoded rRNAs.(Sylvester et al. 2004) Whilst there are no candidate disorders for *MRPL34*, other diseases, such as Parkinson’s disease (previously linked to reduced mtDNA CN (Pyle et al. 2016)), are related to other mitochondrial ribosomal proteins.(Kenmochi et al. 2001; Sylvester et al. 2004; López et al. 2014) This locus also included an additional neighbouring gene, GTP Binding Protein 3 (Mitochondrial) (*GTPBP3*), associated with mtDNA CN at p~5e-07; rare mutations in this gene are known to cause a Leigh syndrome-like disorder.(Kopajtich et al. 2014) However, it is clear that independent replication will be needed in order to confirm these associations, since our current results are in a small sample from our secondary analyses only, and could be attributable to ‘winner’s curse’.(Xiao and Boehnke 2009)

The identification of the cell-count associated locus (*MED24*) could suggest that some loci, such as the chr17 neutrophil count locus, may be related to mtDNA CN only via their association with cell count. It is noteworthy that the meta-analysis in which this locus was identified included ALSPAC mothers with DNA extracted from both white cells and whole blood: it is possible that combining individuals with DNA prepared from multiple sources could lead to preferential detection of loci associated with mtDNA CN predominantly via their association with cellular heterogeneity. When results from participants with more similar DNA sources are pooled, the power to detect loci associated with mtDNA CN *independently* of cell proportions may increase: we postulate that the *BABAM1-ANKLE1-ABHD8-MRPL34* locus might be such a ‘cell-count independent’ locus (although as noted above, this needs further exploration). For loci in this latter category, controlling for cell proportions may improve the signal-to-noise ratio of observed associations, and failure to control for cellular proportions may act akin to measurement error. Future work should seek to assert whether associations of known neutrophil loci with mtDNA CN are entirely due to cell composition in DNA samples, or whether these loci are detected because mtDNA CN is causally related to leucocyte count.(Knez et al. 2016) Ideally, such studies would use directly assayed neutrophil data, as opposed to estimated cell counts. In addition, it might be useful to control associations for platelet count, since platelets are anucleate, but mitochondria-rich.(Urata et al. 2008; Hurtado-Roca et al. 2016)

Beyond the possibility of true underlying genetic heterogeneity between our cohorts, several other technical factors may have limited their comparability. We believe that population stratification is unlikely to be a problem, as analyses controlled for principal components, and the correlation of MAFs for tops hits was high (see **Supplementary Figure 9**). Another technical difference between study groups is the DNA extraction method used. DNA extraction method has a considerable effect on mtDNA CN assay,(Guo et al. 2009) and similar qPCR assays, including that of telomere length.(Raschenberger et al. 2016) However, it is difficult to assess the extent to which DNA extraction method may have affected the performance of the mtDNA CN assay, since extraction method in ALSPAC was also related to age at DNA sampling, and it is possible that the genetic regulation of mtDNA CN varies across the life course.

When we combined all our study groups and looked at whether two hits from a previous *in silico* GWAS were replicated, we found some evidence for one (a SNP in mitochondrial transcription factor A (*TFAM*), but not the other (*CDK6*).(Cai et al. 2015) Despite the comparable sample sizes between our (N=11253) and the previous *in silico* (N=10442 discovery and N=1753 replication) analyses, we observed a considerably smaller effect size for the *TFAM* hit (~1/7 the size of the replication effect from the previous study, despite partial overlap in participants). We postulate that this might flag an important limitation in our work; namely that of measurement error. If a substantial component of our assayed mtDNA CNs includes non-differential measurement error, we would expect to see attenuation of effect sizes in our results. This is possible in qPCR assays: indeed, although we demonstrated correlations in relative mtDNA CNs in a validation analysis, we saw some suggestion of non-linearity. Whilst our attempted replication of the previous *in silico* GWAS hit had 100% power to detect effect sizes of 0.338 SD units (for a SNP with a MAF of 0.169, i.e. the *TFAM* SNP), this power calculation will be overoptimistic if our mtDNA CN assays are affected by measurement error. Therefore, we might suggest that *in silico* measurement of mtDNA CN may have advantages over the directly assayed method in this instance.

In conclusion, we confirm an association of *TFAM* with mtDNA CN, and after performing a range of hypothesis-generating GWAS in diverse study groups, we present several putative regulators of mtDNA CN, that will require further follow-up. However, we generally observed poor concordance across study groups. Overall, our main conclusion is that here we find no strong evidence to support our primary hypothesis of common loci regulating mtDNA CN in the study groups used here. We assess and discuss the possible implications of cellular heterogeneity on our results, and present the directly assayed mtDNA CN assay as another example of a qPCR assay that may be subject to measurement error. These findings should be considered as possible power-limiting factors in GWAS studying the genomic regulation of mtDNA CN. Nonetheless we believe that to fully understand nuclear genomic control of mtDNA CN variation, it is necessary to conduct GWAS of directly assayed mtDNA CN. Thus, our work (the largest GWAS to date) makes an important contribution in terms of future requirements to gain this knowledge, which is necessary for fuller understanding of the biology and potential clinical impact of subtle variation in mtDNA CN.

## Acknowledgements

We are extremely grateful to all the families who took part in ALSPAC, the midwives for their help in recruiting them, and the whole ALSPAC team, which includes interviewers, computer and laboratory technicians, clerical workers, research scientists, volunteers, managers, receptionists and nurses. The UK Medical Research Council and the Wellcome Trust (Grant ref: 102215/2/13/2) and the University of Bristol provide core support for ALSPAC. Child GWAS data was generated by Sample Logistics and Genotyping Facilities at Wellcome Sanger Institute and LabCorp (Laboratory Corporation of America) using support from 23andMe. Mothers GWAS data generated using funding from Wellcome Trust (WT088806).

UKBS used genotype data from cases and population controls that were generated by the Wellcome Trust Case Control Consortium 2 (PMID: 17554300, a full list of the investigators who contributed to the generation of the data is available from http://www.wtccc.org.uk). The UK Blood Services Collection of Common Controls were funded by The Wellcome Trust.

This research was funded by a Medical Research Council (MRC) grant awarded to Santiago Rodriguez (MR/K002767/1). AG is funded by a Wellcome Trust PhD studentship (102433/Z/13/Z). RB is in receipt of a Wellcome Trust Centre for Mitochondrial Research PhD Studentship (Grant ref: 096919/Z/11/Z), as well as funding from The Barbour Foundation. GH is a PDUK Senior Research Fellow (Grant ref: F-1202) and receives funding from the Wellcome Trust Centre for Mitochondrial Research at Newcastle University (Grant ref: 096919/Z/11/Z). Work was carried out in the MRC Integrative Epidemiology Unit at the University of Bristol (MC_UU_12013/5 and MC_UU_12013/8), and at the Wellcome Trust Centre for Mitochondrial Research at Newcastle University. Additional support for study recruitment and sample collection was provided by the British Heart Foundation (SP/07/008/24066), UK Medical Research Council (G1001357), and Wellcome (WT092830M and WT088806). PFC is a Wellcome Trust Senior Fellow in Clinical Science (101876/Z/13/Z), and a UK NIHR Senior Investigator, who receives support from the Medical Research Council Mitochondrial Biology Unit (MC_UP_1501/2), the Medical Research Council (UK) Centre for Translational Muscle Disease (G0601943), and the National Institute for Health Research (NIHR) Biomedical Research Centre based at Cambridge University Hospitals NHS Foundation Trust and the University of Cambridge. The views expressed are those of the author(s) and not necessarily those of the NHS, the NIHR or the Department of Health. DAL is a UK National Health Research Senior Investigator (NF-SI-0611-10196). This publication is the work of the authors, who will serve as guarantors for the contents of this paper.

## Notes

**Conflict of Interest**: TRG reports funding from Sanofi, Biogen, and GlaxoSmithKline for projects unrelated to the work presented in this manuscript.

